# The CIC-DUX4 fusion oncoprotein drives metastasis and tumor growth via distinct downstream regulatory programs and therapeutic targets in sarcoma

**DOI:** 10.1101/476283

**Authors:** Ross A. Okimoto, Wei Wu, Shigeki Nanjo, Victor Olivas, Yone K. Lin, Rieko Oyama, Tadashi Kondo, Trever G. Bivona

**Affiliations:** Department of Medicine, University of California, San Francisco, San Francisco, California, USA; Helen Diller Family Comprehensive Cancer Center, University of California, San Francisco, San Francisco, California, USA; Division of Rare Cancer Research, National Cancer Center Research Institute, Tokyo, Japan; Cellular and Molecular Pharmacology, University of California, San Francisco, San Francisco, California, USA

## Abstract

Transcription factor fusion genes create oncoproteins that drive oncogenesis, and represent challenging therapeutic targets. Understanding the molecular targets by which such fusion oncoproteins promote malignancy offers an approach to develop rational treatment strategies to improve clinical outcomes. CIC-DUX4 is a transcription factor fusion that defines certain undifferentiated round cell sarcomas with high metastatic propensity and poor clinical outcomes. The molecular targets regulated by the CIC-DUX4 oncoprotein that promote this aggressive malignancy remain largely unknown. We show that increased expression of *ETV4* and *CCNE1* occurs via neo-morphic, direct effects of CIC-DUX4 and drives tumor metastasis and survival, respectively. We demonstrate a molecular dependence on the CCNE-CDK2 cell cycle complex that renders CIC-DUX4 tumors sensitive to inhibition of the CCNE-CDK2 complex, highlighting a therapeutic strategy for CIC-DUX4 tumors. Our findings highlight a paradigm of functional diversification of transcriptional repertoires controlled by a genetically-aberrant transcriptional regulator, with therapeutic implications.

## Introduction

The biological regulation of transcription factors and repressor proteins is an essential mechanism to maintain cellular homeostasis, and is often dysregulated in human cancer ^1^. Indeed, chromosomal rearrangements involving transcriptional regulatory genes that lead to transcriptional dysregulation are present in many cancers, including ~30% of all soft-tissue sarcomas ^2^. The majority of these oncogenic fusions involve transcription factors or regulators that are not readily “druggable” in a direct pharmacologic manner, and thus have proven difficult to therapeutically target in the clinic. A prime example is the CIC-DUX4 fusion oncoprotein, which fuses *Capicua* (*CIC*) to the double homeobox 4 gene, *DUX4*. CIC-DUX4-positive soft-tissue tumors are an aggressive subset of undifferentiated round cell sarcomas that arise in children and young adults. Despite histological similarities to Ewing sarcoma, CIC-DUX4-positive sarcomas are clinically-distinct and typically characterized by the rapid development of lethal metastatic disease and chemo-resistance ^3^. The unique cytogenetic and clinical features that distinguish CIC-DUX4-positive sarcomas from other small round cell tumors provides an opportunity for specific precision medicine-based therapies to improve clinical outcomes.

CIC is a transcriptional repressor protein ^4^. The CIC-DUX4 fusion structurally retains >90% of native CIC, yet it functions as a transcriptional activator instead of a transcriptional repressor ^5,6^. This property suggests that the C-terminal DUX4 binding partner may confer neo-morphic transcriptional regulatory properties to CIC, while retaining wild-type CIC DNA binding specificity. *ETV4* is one of the most well-characterized transcriptional targets of CIC-DUX4 ^5,7,8^. Over 90% of CIC-DUX4 tumors show ETV4 upregulation by immunohistochemistry (IHC), which distinguishes them from other small round blue cell sarcomas ^8^. We and others have demonstrated a pro-metastatic function for ETV4 overexpression in tumors with inactivation of wild-type (WT) CIC ^9,10^. The functional role of ETV4 in CIC-DUX-positive tumors is unknown.

Beyond ETV4, the identity and function of other CIC target genes are less well-defined. Intriguingly, recent studies showed increased expression of cell-cycle regulatory genes in CIC-rearranged sarcomas, although the functional relevance of this observation for oncogenesis and cancer growth in this context is unclear ^6,7^.

Here, we develop a range of *in vitro* and *in vivo* cancer model systems to define the mechanism by which CIC-DUX4 co-opts transcriptional pathways that native CIC controls to promote hallmark features of malignancy, including tumor cell survival, growth, and metastasis.

## Results and Discussion

We recently reported that inactivation of native CIC de-represses an ETV4-MMP24 mediated pro-metastatic circuit ^9^. Since *ETV4* is a direct transcriptional target of CIC-DUX4 (Fig. S1A) ^6,7^, we hypothesized that the high metastatic rate observed in patients harboring CIC-DUX4 fusions was dependent on ETV4 expression. To explore this hypothesis, we developed an orthotopic soft-tissue metastasis model that utilizes luciferase-based imaging to track tumor dissemination *in vivo* (Fig. 1A). This system produces rapid pulmonary metastases that accurately recapitulates metastatic tumor dissemination in human patients ^3^. Using this *in vivo* system, we engineered NIH-3T3 mouse fibroblasts that offer the advantage of a genetically-controlled system (as with the study of other oncoproteins), to express the CIC-DUX4 fusion oncoprotein^11–13^. We observed rapid primary tumor formation at the site of implantation in 100% of the injected mice (Fig. 1B). Genetic silencing of *ETV4* decreased expression of its established target MMP24 ^9^ (Fig. 1C), and significantly impaired metastatic efficiency *in vivo* and invasive capacity of cells *in vitro*, compared to mice bearing a control silencing vector (Fig. 1B, 1D, and 1E). While ETV4 suppression decreased distant pulmonary metastases, it did not have a profound impact on tumor growth when CIC-DUX4 expressing cells were implanted either orthotopically or subcutaneously into the flank of immunocompromised mice (Fig. 1F-G). These findings suggest that the primary function of CIC-DUX4-mediated *ETV4* upregulation is to promote invasion and metastasis, but not tumor cell proliferation or tumor growth *per se*.

**Figure 1.**
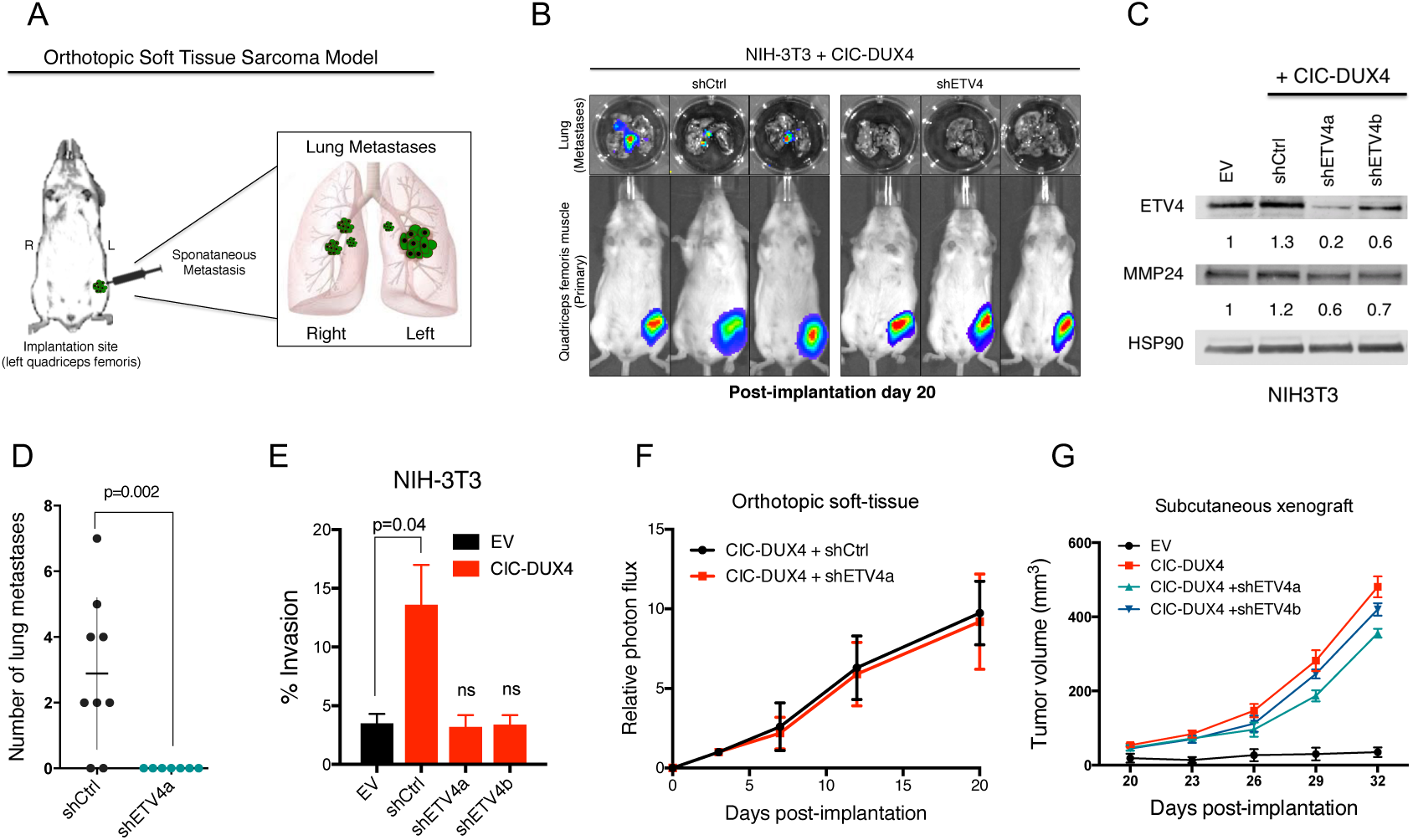
ETV4 promotes metastasis in CIC-DUX4 sarcoma. A) Schematic of the orthotopic soft-tissue metastasis model. B) Representative bioluminscent (BLI) images from mice bearing CIC-DUX4 NIH-3T3 cells with shCtrl (n=9) or shETV4 (n=7). C) Immunoblot of ETV4 and MMP24 from NIH3T3 cells expressing EV, shCtrl, shETV4a, or shETV4b. D) Number of lung metastases in mice bearing CIC-DUX4 NIH-3T3 cells with shCtrl or shETV4. E) Transwell invasion assay comparing CIC-DUX4 expressing NIH-3T3 cells with either EV, shCtrl, shETV4a, or shETV4b. F) Relative photon flux from mice orthotopically implanted with CIC-DUX4 NIH-3T3 cells expressing either shCtrl or shETV4. G) Subcutaneously implanted NIH-3T3 cells with either EV, CIC-DUX4, or CIC-DUX4 with shETV4a or shETVb.

We next investigated transcriptional targets and programs that could regulate other key aspects of tumor biology beyond metastasis, including tumor cell proliferation and growth. Prior studies suggested that the CIC-DUX4 fusion can regulate cell-cycle gene expression ^6,7^. We established that ectopic expression of CIC-DUX4 in NIH-3T3 cells increased tumor growth *in vitro* and *in vivo* through enhanced cell-cycle progression (Fig. S1B-S1E). The CIC-DUX4 fusion increased the number of cells progressing through S-phase, as reflected by an increased G2/M fraction compared to control (Fig. S1F-G). This CIC-DUX4 mediated cell-cycle progression was reversed upon 5-fluorouracil (5-FU) treatment (Fig. S1F-G). Since 5-FU induces a G1/S arrest, we hypothesized that the CIC-DUX4 fusion regulates late G1 and S phase-promoting genes. In order to identify these CIC-DUX4 regulated cell-cycle genes, we leveraged a publicly available microarray–based dataset (GSE60740) to perform a comparative transcriptional analysis between CIC-DUX4 replete (IB120 EV cell line) and two independent CIC-DUX4 knockdown (KD) patient-derived cell lines (IB120 shCIC-DUX4a and IB120 shCIC-DUX4b) ^14^. Upon CIC-DUX4 KD, we observed 409 and 205 down-regulated genes (logFC <^-^2, FDR <0.05) in IB120 shCICDUX4a and IB120 shCIC-DUX4b, respectively (Fig. 2A). There were 165 shared genes between the two shCIC-DUX4 datasets. Functional clustering of the 165 putative CIC-DUX4 target genes revealed enrichment for genes that regulate DNA replication and cell-cycle machinery (Fig. 2B). By comparing the differential expression of these 165 genes between 14 CIC-DUX4 and 6 EWS-NFATC2 patient-derived tumors (GSE60740), we observed a significant increase in expression of multiple cell-cycle regulatory genes in CIC-DUX4 tumors (Fig. 2C). Our findings extend recent studies^6,7^, and indicate that targeting the cell-cycle in CIC-DUX4 tumors is a potential therapeutic approach.

**Figure 2.**
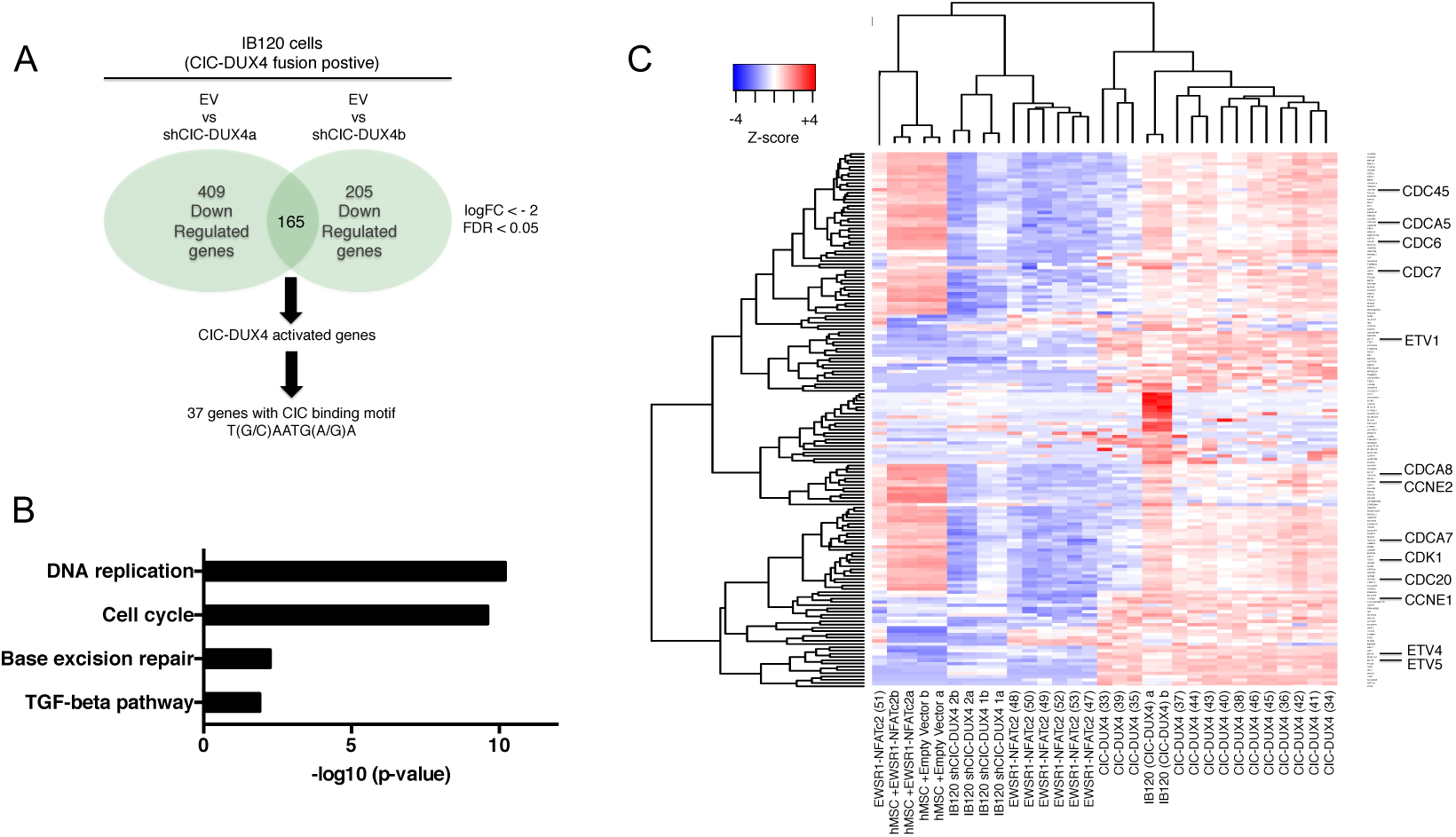
CIC-DUX4 regulates cell-cycle genes. A) Schematic algorithm to identify putative CIC-DUX4 target genes. B) Functional annotation of CIC-DUX4 activated genes in IB120 cells. C) Heatmap depicting the 165 CIC-DUX4 activated genes identified in Fig. 2A across 14 CIC-DUX4 and 7 EWSR1-NFATc2 patient derived tumors. Cell-cycle genes are magnified.

In order to identify direct transcriptional targets of the CIC-DUX4 fusion, we first surveyed the promoters (within −2kb and +150bp of the transcriptional start site) of all 165 putative response genes for the highly conserved CIC binding motif (T(G/C)AATG(A/G)A)^5^. Using this systematic approach, we identified 37 genes, including the known CIC target genes ETV1/4/5 and multiple regulators of the cell-cycle (Table S1). Since ectopic expression of CIC-DUX4 promoted S-phase progression (Fig. S1F-G), we focused on genes that directly regulated the G1/S transition. Of these genes, *CCNE1* expression was consistently upregulated in CIC-DUX4 tumors (Fig. 2C). We therefore investigated whether CIC-DUX4 transcriptionally controls *CCNE1* expression. To explore this hypothesis, we first localized two tandem CIC-binding motifs within 1kb of the *CCNE1* transcriptional start site (Fig. 3A). We next performed chromatin immunoprecipitation PCR (ChIP-PCR) analysis, which revealed *CCNE1* promoter occupancy by the CIC-DUX4 fusion (Fig. 3B-C). Additionally, a luciferase-based reporter assay demonstrated enhanced promoter activity by the ectopic expression of CIC-DUX4 compared to CIC wild-type and empty-vector (EV) control (Fig. 3D). These data show that *CCNE1* is a direct transcriptional target that is upregulated by CIC-DUX4.

**Figure 3.**
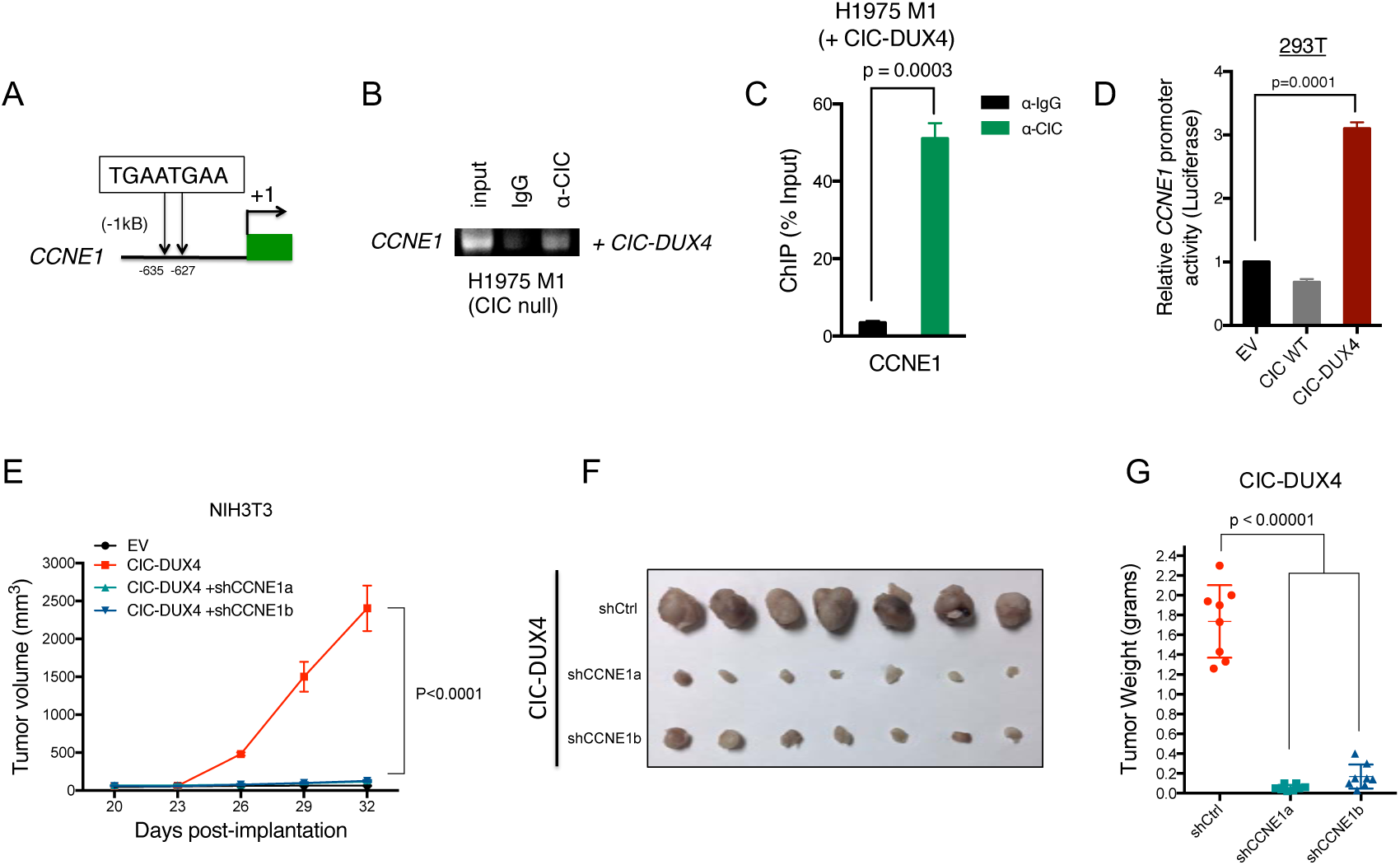
CCNE1 is a molecular target of CIC-DUX4. A) Schematic of two CIC biding motifs in the *CCNE1* promoter. B-C) ChIP-PCR from H1975 M1 (CIC wild-type null) cells reconstituted with CIC-DUX4 showing CIC-DUX4 occupancy on the *CCNE1* promoter. D) *CCNE1* luciferase promoter assay in 293T cells comparing EV, CIC wild-type, or CIC-DUX4. E) Subcutaneously implanted NIH-3T3 cells expressing either EV, CIC-DUX4, or CIC-DUX4 with shCCNE1a or shCCNE1b. F) Tumor explants from mice in 3E. G) Tumor weights from mice in 3E.

To explore the functional role of CCNE1 in CIC-DUX4-expressing tumors, we genetically-silenced *CCNE1* in CIC-DUX4 expressing NIH-3T3 cells (a genetically-controlled system) and observed decreased tumor growth *in vitro* and *in vivo* compared to control (Fig. 3E-G and S2A-C). While there is no validated pharmacologic strategy to directly block CCNE1, inhibition of the CCNE1 binding partner CDK2 has therapeutic efficacy in other cancer types ^15–17^. We hypothesized that inhibiting CDK2 in CIC-DUX4 expressing tumors with the established small molecule drug dinaciclib^15^ could limit tumor growth. To explore this hypothesis, we implanted CIC-DUX4 expressing NIH-3T3 cells subcutaneously into immune-deficient mice and treated with low-dose dinaciclib (20mg/kg/day) and observed decreased tumor growth compared to vehicle control (Fig. 4A-C) ^15^. These findings suggest that pharmacologic inhibition of the CCNE-CDK2 complex is a potential therapeutic strategy in CIC-DUX4 expressing tumors. There are few patient-derived models of CIC-DUX4-positive tumors. Nevertheless, in order to increase the clinical relevance of our findings we obtained rare, established patient-derived CIC-DUX4 expressing cells (NCC-CDS1-X1 and NCC-CDS-X3) ^18^ to test the functional impact of *ETV4* KD and CDK2 inhibition. Consistent with the findings arising from our isogenic NIH-3T3 system, we found that genetic silencing of ETV4 decreased invasiveness, but did not impact tumor growth (Fig. S3A-B). Additionally, we found that CIC-DUX4 expressing cells were exquisitely sensitive to nanomolar (nM) concentrations of two established, independent CDK2 inhibitors, dinaciclib and SNS-032 ^15,19^ (Fig. 4D, S3C-E). The effects on tumor viability were largely mediated through apoptotic cell death as measured by caspase activity and PARP cleavage, again indicating that CIC-DUX4 tumors are dependent on the CCNE-CDK2 complex for survival (Fig. 4E-G).

**Figure 4.**
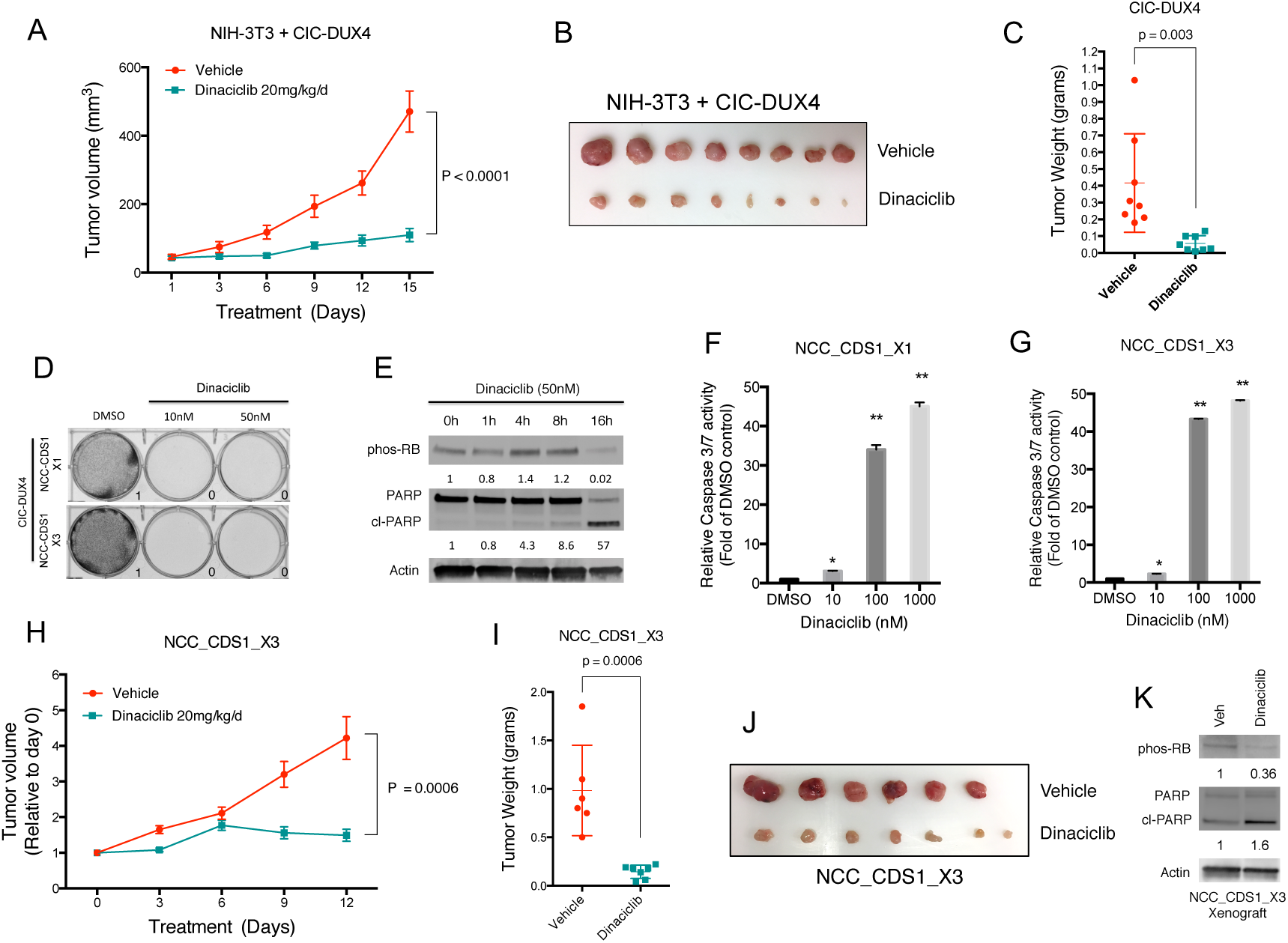
CCNE-CDK2 complex is a therapeutic target in CIC-DUX4 sarcoma. A) Subcutaneously implanted NIH-3T3 cells expressing CIC-DUX4 and treated with vehicle or dinaciclib. B) Tumor explants from mice in 4A. C) Tumor weights from mice in 4A. D) Patient derived CIC-DUX4 expressing cells (NCC_CDS1_X1 and NCC_CDS_X3) treated with dinaciclib or DMSO. E) Immunoblot of phosphorylated-Rb, PARP, and actin control in NCC_CDS1_X3 cells treated with dinaciclib. F) Relative caspase 3/7 activity in NCC_CDS1_X1 and; G) NCC_CDS_X3 cells treated with dinaciclib or DMSO. **p-value < 0.0001. H) Subcutaneously implanted NCC_CDS1_X3 cells treated with vehicle control or dinaciclib. I) Tumor weights from mice treated in 4H. J) Tumor explants from mice in 4H. K) Immunoblot of phosphorylated-RB, total and cleaved PARP, and actin control from a NCC_CDS1_X3 tumor explant treated with vehicle or dinaciclib.

We next tested whether CDK2 inhibition could specifically suppress the growth of patient-derived CIC-DUX4-expressing cells *in vivo*. To address this hypothesis, we generated subcutaneous tumor xenografts of the patient-derived CIC-DUX4 expressing cells in immune-deficient mice and found that dinaciclib limited tumor growth compared to vehicle control treatment (Fig. 4H-J). The decreased tumor growth observed in dinaciclib-treated mice was accompanied by increased PARP cleavage in tumor explants, consistent with the apoptotic effect observed *in vitro* (Fig. 4K). The impact on tumor growth following CDK2 inhibition was relatively specific for CIC-DUX4 tumors, as we did not observe similar therapeutic responses to dinaciclib in other sarcoma subtypes that harbor distinct transcriptional factor fusion oncoproteins (Rhabdomyosarcoma or Ewing Sarcoma), *in vitro* or *in vivo* (Fig. S4A-B). Consistent with these findings, we observed low levels of *CCNE1* in PAX3-FOXO1-positive RH30 rhabdomyosarcoma cells compared to CIC-DUX4 expressing NCC-CDS-X1 cells (Fig. S4C). These findings further suggest that the CCNE-CDK2 complex is a specific molecular dependence and therapeutic target in CIC-DUX4-expressing tumors.

To further demonstrate the therapeutic specificity of targeting CDK2, we used the clinical CDK4/6 inhibitor palbociclib that has no substantial activity against CDK2, and found no impact on tumor growth or apoptosis in NCC-CDS-X1 cells (Fig. S5A). Moreover, while genetic silencing of *CDK2* with siRNAs did not impact cell invasion, it did decrease cell number and viability in CIC-DUX4-expressing NCC-CDS-X1 cells compared to control (Fig. S5B-D). While we did not observe an effect on viability with *CCNE1* knockdown (KD) alone, combined *CCNE1* and *CCNE2* KD reduced viability to similar levels as si*CDK2* cells (Fig. S5C-D). These findings are consistent with prior observations that both CCNE1 and CCNE2 converge on, and activate CDK2, which our pharmacologic and genetics studies described above indicate is required for CIC-DUX4-expressing tumor survival^20^.

Our data suggest that in patient-derived NCC-CDS-X1 cells CCNE2 can compensate for CCNE1 loss. Consistent with this finding, we observed significant (~100-fold) upregulation of *CCNE2* mRNA upon genetic silencing of *CCNE1* (Fig. S5F) in NCC-CDS-X1 cells. In contrast to these findings, we did not observe a compensatory increase in *CCNE2* expression with *CCNE1* KD in NIH-3T3 (mouse) cells expressing CIC-DUX4 (Fig. S5G). The lack of *CCNE2* upregulation in CIC-DUX4-expressing NIH-3T3 cells is a plausible explanation for the growth suppressive effect of *CCNE1* KD in our initial NIH-3T3 system (Fig. 3E), a phenotype that was not shared with patient-derived NCC-CDS cells (Fig. S5B). Collectively, our data suggest that human CIC-DUX4-expressing cells have a specific dependence on the CCNE-CDK2 cell-cycle regulatory complex. Our mechanistic dissection of CIC-DUX4 tumors reveals a unique molecular and therapeutic dependence on the CCNE-CDK2 cell cycle complex. Our data show that this dependence can be exploited with clinical CDK2 inhibitors that limit tumor growth through apoptotic induction in a tumor type with few effective therapies.

The CIC-DUX4 fusion oncoprotein is a relatively understudied molecular entity that characterizes a rare, albeit lethal subset of undifferentiated round cell sarcomas. We undertook a mechanistic dissection of the molecular function of the CIC-DUX4 fusion oncoprotein and report unique dependencies on specific transcriptional targets of the fusion oncoprotein that promote either tumor growth or metastasis. We reveal that *ETV4* is a conserved target gene that enhances the metastatic capacity of CIC-DUX4 tumors, without significantly impacting tumor growth. These findings extend previous data that demonstrate a pro-metastatic role for ETV4 in certain human cancer s^9,21^. Targeting an ETV4-mediated transcriptional program downstream of CIC-DUX4 can potentially limit tumor dissemination.

Further analysis of CIC-DUX4-bearing tumors coupled with our functional studies also identified a dependence on specific cell-cycle machinery to enhance tumor growth, but not invasive capacity. We reveal that the CCNE-CDK2 complex is a molecular target of the CIC-DUX4 oncoprotein that controls tumor growth and survival. While others have observed transcriptional upregulation of cell-cycle genes in CIC-DUX4 tumors^6,7^, our data establish the CCNE-CDK2 complex as a direct molecular target that can be pharmacologically exploited with clinically-developed CDK2 inhibitors. The therapeutic impact may extend beyond CIC-DUX4 fusion oncoproteins, to include other CIC-fused oncoproteins. Recent findings reveal a shared transcriptional program downstream of all known CIC fusions, including CIC-FOXO4 and CIC-NUTM1^22^. These findings suggest that many CIC-fused tumors retain their CIC binding specificity while converting native CIC into a transcriptional activator instead of a repressor in a neo-morphic functional manner. It would be compelling to explore if the CCNE-CDK2 complex or other components of the cell-cycle machinery drive tumor growth in these other CIC-fused tumor types, an area for future investigation.

Our mechanistic dissection of the downstream molecular targets of CIC-DUX4 provides therapeutically-relevant insight for targeting ETV4-mediated metastasis and CCNE-CDK2-regulated tumor growth. Our data enforce a broader conceptual paradigm in which certain transcription factor fusion oncoproteins, such as CIC-DUX4, utilize neo-morphic and distinct downstream regulatory programs to control divergent cancer hallmarks, such as proliferative capacity and metastatic competency. The data offer a potential mechanistic explanation for the pleiotropic functions of this important class of oncoproteins (i.e. transcription factor fusion oncoproteins such as CIC-DUX4).

Our findings highlight the clinical importance of molecular sub-classification of morphologically-similar tumor types, such as small round cell sarcomas. Identifying the different fusion oncoproteins present in clinical samples paves the way for oncoprotein fusion-specific therapeutic targeting to improve patient outcomes. Our study highlights the utility of elucidating the mechanistic features of tumors that are driven by transcription factor fusion oncoproteins to identify precision medicine-based, molecular therapeutic strategies.

## Methods

### Orthotopic and subcutaneous soft tissue xenografts in immunodeficient mice

Six to eight-week old female SCID mice were purchased from Taconic (Germantown, NY). Specific pathogen-free conditions and facilities were approved by the American Association for Accreditation of Laboratory Animal Care. Surgical procedures were reviewed and approved by the UCSF Institutional Animal Care and Use Committee (IACUC), protocol #AN107889-03A.

To prepare cell suspensions for quadriceps injection, adherent tumor cells were briefly trypsinized, quenched with 10% FBS RPMI media and resuspended in PBS. Cells were pelleted again and mixed with Matrigel matrix (BD Bioscience 356237) on ice for a final concentration of 1.0×10^5^ cells/μl. The Matrigel-cell suspension was transferred into a 1ml syringe and remained on ice until the time of implantation.

For orthotopic injection, mice were placed in the right lateral decubitus position and anesthetized with 2.5% inhaled isoflurane. A 0.5 cm surgical incision was made along the posterior medial line of the left hindlimb, fascia and adipose tissue layers were dissected and retracted to expose the quadriceps femoris muscle. A 30-guage hypodermic needle was used to advance through the muscular capsule. For all cell lines, care was taken to inject 10μl (1.0×10^6^ cells) of cell suspension directly into the left quadriceps femoris. The needle was rapidly withdrawn and mice were observed for bleeding. Visorb 4/0 polyglycolic acid sutures were used for primary wound closure of the fascia and skin layer. Mice were observed post-procedure for 1-2 hours and body weights and wound healing were monitoring weekly. For PDX subcutaneous xenotransplantation, 3.0×10^6^ NCC_CDS1_X3 cells were resuspended in 50% PBS/50% Matrigel matrix and injected into the flanks of immunodeficient mice.

### In-vivo bioluminescence imaging

Mice were imaged at the UCSF Preclinical Therapeutics Core starting on post-injection day 7 with a Xenogen IVIS 100 bioluminescent imaging system. Prior to imaging, mice were anesthetized with isoflurane and intraperitoneal injection (IP) of 200μl of D-Luciferin at a dose of 150mg/kg body weight was administered. Weekly monitoring of bioluminescence of the engrafted hindlimb tumors was performed until week 5. Radiance was calculated automatically using Living Image Software following demarcation of the thoracic cavity (ROI) in the supine position. The radiance unit of photons/sec/cm2/sr is the number of photons per second that leave a square centimeter of tissue and radiate into a solid angle of one steradian (sr).

### Ex-vivo bioluminescence imaging

Mice were injected IP with 200 μl (150mg/kg) of D-Luciferin and subsequently sacrificed at 5 weeks, en-bloc resection of the heart and lungs was performed. The heart was removed and the lungs were independently imaged. Imaging was performed in a 12 well tissue culture plate with Xenogen IVIS 100 bioluminescent imaging.

### Cell lines and culture reagents

Cell lines were cultured as recommended by the American Type Culture Collection (ATCC). NIH-3T3, 293T, A673, SAOS2 cells were obtained ATCC. NCC_CDS1_X1 and NCC_CDS_X3 were obtained from Tadashi Kondo at the National Cancer Center, Tokyo, Japan. The presence of the CIC-DUX4 fusion was confirmed through RNAseq analysis using the “grep” command as previously described (Panagopoulos et al, Plos One 2014). All cell lines were maintained at 37 °C in a humidified atmosphere at 5% CO_2_ and grown RPMI 1640 media supplemented with 10% FBS, 100 IU/ml penicillin and 100ug/ml streptomycin.

Dinaciclib, palbociclib, SNS-032 was purchased from SelleckChem.

### Gene knockdown and over-expression assays

All shRNAs were obtained from Sigma Aldrich. Sequences for individual shRNAs are as follows:

shETV4a: catalog # TRCN0000055132.

shETV4b: catalog # TRCN0000295522.

shCCNE1a: catalog # TRCN0000222722

shCCNE1b: catalog # TRCN0000077776

shCCNE1b: catalog # TRCN0000077777

ON-TARGET plus ETV4, Scramble, CDK2, CDK1, CCNE1, CCE2 siRNA was obtained from GE Dharmacon and transfection performed with Dharmafect transfection reagent. The HA-tagged CIC-DUX4 plasmid was obtained from Takuro Nakarmua, Tokyo, Japan. Sequence verification was performed using sander sequencing. The lentiviral GFP-Luciferase vector was a kind gift from Michael Jensen (Seattle Children’s Research Institute, Seattle). Fugene 6 transfection reagent was used for all virus production and infection was carried out with polybrene.

### Chromatin immunoprecipitation and PCR

CIC null cells (H1975 M1) were transfected with either GFP control, wild-type CIC, or CIC-DUX4 for 48 hours. SimpleCHIP Enzymatic Chromatin IP Kit (Cell Signaling Technology) was used with IgG (Cell signaling Technology) and CIC (Acris) antibodies per the manufactures protocol.

ETV4 PCR primers were previously described (Okimoto et al., Nature Genetics 2016). The ETV4 primer sequences were as follows:

ETV4_Foward 5’-CGCATCAGACCCAAGACCGTGG-3’

ETV4_Reverse 5’-CCGGAGAGTCGTCCGGCCTGG-3’

CCNE1 PCR primers were designed to flank a tandem TGAATGAA/TGAATGAA sequence from positions −635 to −628 and −627 to −620 in the CCNE1 promoter. The primer sequences were as follows:

CCNE1_1F CGTCTCGGCCTCCCACAATGCTGGG and CCNE1_1R CGCGCCTGTGCCTTGGCCTAGAACC.

### Luciferase promoter assay

293T cells were obtained from ATCC. Cells were grown in Dulbecco’s modified Eagle Medium (DMEM), supplemented with 10% FBS, 100 IU/ml penicillin and 100ug/ml streptomycin in a 5% CO_2_ atmosphere. Cells were split into a 96 well plate to achieve 50% confluence the day of transfection. LightSwitch luciferase assay system (SwitchGear Genomics S720355) was used per the manufactures protocol. Briefly, a mixture containing FuGENE 6 transfection reagent, 50ng Luciferase GoClone *CCNE1* promoter (#S720355) plasmid DNA, 50ng of either control (empty) vector or CIC-DUX4 or wild-type CIC was added to each well. All transfections were performed in quintuplicate.

### Western blot and qRT-PCR

All immunoblots represent at least two independent experiments. Adherent cells were washed and lysed with RIPA buffer supplemented with proteinase and phosphatase inhibitors. Proteins were separated by SDS-PAGE, transferred to Nitrocellulose membranes, and blotted with antibodies recognizing: GFP (Cell Signaling), E-Cadherin (Cell Signaling), HSP90 (Cell Signaling), ETV4 (Lifespan), CCNE1 (Cell Signaling), PARP (Cell Signaling), Phosphor-RB (Cell Signaling), Actin (Sigma), HA-tag (Cell Signaling).

### Xenograft tumors

Subcutaneous xenografts were explanted on day 4 of treatment. Tumor explants were immediately immersed in liquid nitrogen and stored at −80 degrees. Tumors were disrupted with a mortar and pestle, followed by sonication in RIPA buffer supplemented with proteinase and phosphatase inhibitors. Proteins were separated as above. Antibodies to PARP and phosphor-RB were both from Cell Signaling.

Isolation and purification of RNA was performed using RNeasy Mini Kit (Qiagen). 500 ng of total RNA was used in a reverse transcriptase reaction with the SuperScript III first-strand synthesis system (Invitrogen). Quantitative PCR included four replicates per cDNA sample. Human (*CDK1, CDK2, CCNE1, CCNE2, GAPDH*, and *TBP)* and mouse (*CCNE1*, *CCNE2*, and *GAPDH*) were amplified with Taqman gene expression assays (Applied Biosystems). Expression data was acquired using an ABI Prism 7900HT Sequence Detection System (Applied Biosystems). Expression of each target was calculated using the 2^−ΔΔCt^ method and expressed as a relative mRNA expression.

### Transwell migration and invasion assays

RPMI with 10% FBS was added to the bottom well of a trans-well chamber. 2.5×10^4^ cells resuspended in serum free media was then added to the top 8 μM pore matrigel coated (invasion) or non-coated (migration) trans-well insert (BD Biosciences). After 20 hours, non-invading cells on the apical side of inserts were scraped off and the trans-well membrane was fixed in methanol for 15 minutes and stained with Crystal Violet for 30 minutes. The basolateral surface of the membrane was visualized with a Zeiss Axioplan II immunofluorescent microscope at 10X. Each trans-well insert was imaged in five distinct regions at 10X and performed in triplicate. % invasion was calculated by dividing the mean # of cells invading through Matrigel membrane / mean # of cells migrating through control insert.

### Immunostaining (IHC): subcutaneous xenografts

Formalin fixed, paraffin embedded (FFPE) patient derived tumor specimens were obtained under the auspices of institutional review board (IRB)-approved clinical protocols. Specimens were de-paraffinized and stained with an antibody against Cleaved Caspase 3 (Cell Signaling) and phosphorylated RB (Cell Signaling).

### Establishment of a CIC responsive gene set and identification of CIC-DUX4 target genes

A publically curated Affymetrix mRNA dataset (GSE60740) of 14 CIC-DUX4, 7 EWSR1-NFATc2 tumors, and a CIC-DUX4 expressing cell line (IB120) expressing either shRNA’s targeting CIC-DUX4 or control was used to generate a list of CIC-DUX4 responsive genes. Specifically, we first independently compared IB120 cells expressing either EV control to each individual shRNA targeting the CIC-DUX4 fusion (shCIC-DUX4a and shCICDUX4b). Using logFC <-2 and FDR<0.05 to identify the most down-regulated genes, we then generated a shared gene list (N = 165) that we referred to as “CIC-DUX4 responsive genes”. We then used the CIC-DUX4 responsive gene set to perform functional clustering with Database for Annotation, Visualization, and Integrated Discovery (DAVID).

Using the CIC-DUX4 responsive gene set, we generated a gene expression heat map comparing: 1) IB120 cells expressing control vector and; 2) the two independent shRNAs targeting CIC-DUX4; 3) the 14 CIC-DUX4 patient derived tumors and; 4) the 7 EWSR1-NFATc2 tumors. Hierarchical clustering was performed using the differentially expressed CIC-DUX4 responsive gene set. We performed a similar hierarchical comparison of PAX3-FOXO1 positive cell lines (RH30) to CIC-DUX4 positive NCC-CDS-X1 cells as documented above.

To identify putative CIC-DUX4 target genes, we surveyed all 165 CIC responsive genes for the CIC-binding motif (TGAATGAA) within −2000bp and +150bp of the transcriptional start site. 37 of the 165 genes contained the CIC-binding motif (supplementary table). Promoter sequences were download from eukaryotic promoter database (http://epd.vital-it.ch/).

### Cell cycle analysis

To determine the effect of CIC-DUX4 expression on cell cycle, NIH-3T3 cell lines were cultured to ~70% confluence and transfected with CIC-DUX4 or a GFP control vector for 48 hours. Cells were trypsinized and fixed in ice cold ethanol for 10 minutes and subsequently stained with propidium iodide (PI) solution (Sigma Aldrich) at room temperature for 15 minutes. Cells were analyzed on a BD LSRII flow cytometer.

### Statistical analysis

Experimental data are presented as mean +/-SEM. P-values derived for all in-vitro experiments were calculated using two-tailed Student’s t test.

## Supporting information

## Author contributions

R.A.O. designed and performed the experiments, analyzed the data, and wrote the manuscript. R.A.O., V.O., S.N., R. O. and T.K. performed mouse studies. W.W. performed bioinformatics. T.K. performed experiments and provided cell lines. T.G.B. directed the project, analyzed experiments, and wrote the manuscript.

## Acknowledgements

R.A.O. was supported by a grant from the A.P. Giannini Foundation and 1K08CA222625-01. T.G.B acknowledges support from NIH / NCI U01CA217882, NIH / NCI U54CA224081, NIH / NCI R01CA204302, NIH / NCI R01CA211052, NIH / NCI R01CA169338, and the Pew-Stewart Foundations. The authors thank Dr. Takuro Nakamura for providing the HA-tagged CIC-DUX4 plasmid.

