## Supplementary material for "The CIC-DUX4 fusion oncoprotein drives metastasis and tumor growth via distinct downstream regulatory programs and therapeutic targets in sarcoma"

Supplementary data contains 5 figures and 1 table.

Supplemental Figure 1. CIC-DUX4 regulates cell-cycle progression and tumor growth.

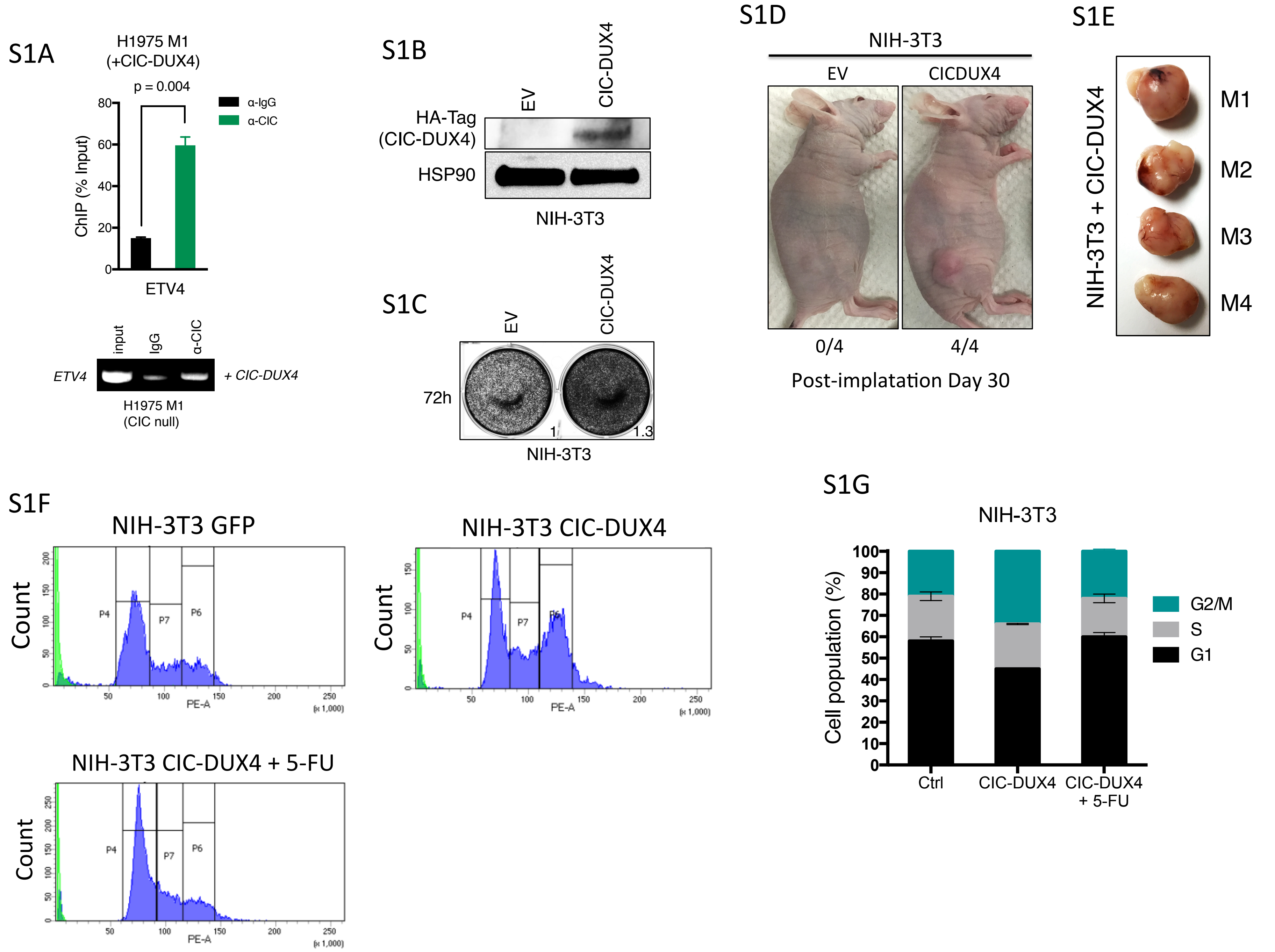

**Supplemental Figure 1. CIC-DUX4 regulates cell-cycle progression and tumor growth.** A) ChIP-PCR from H1975 M1 (CIC wild-type null) cells reconstituted with CIC-DUX4 showing CIC-DUX4 occupancy on the *ETV4* promoter. B) Immunoblot of CIC-DUX4 (HA-tag) and HSP90 in NIH-3T3 cells. C) Crystal violet assay comparing NIH-3T3 cells expressing either EV control or CIC-DUX4. D) Subcutaneously implanted NIH-3T3 cells expressing either EV control (n=4) or CIC-DUX4 (n=4). E) Tumor explants from mice in S1C. F) Cell-cycle profiles of NIH-3T3 cells expressing GFP control, CIC-DUX4, or CIC-DUX4 treated with 5-fluorouracil. G) Cell-cycle distribution of NIH-3T3 cells expressing either EV control, CIC-DUX4 alone or with 5-fluorouracil.

Supplemental Figure 2. CCNE1 inhibition decreases tumor growth in CIC-DUX4 expressing cells.

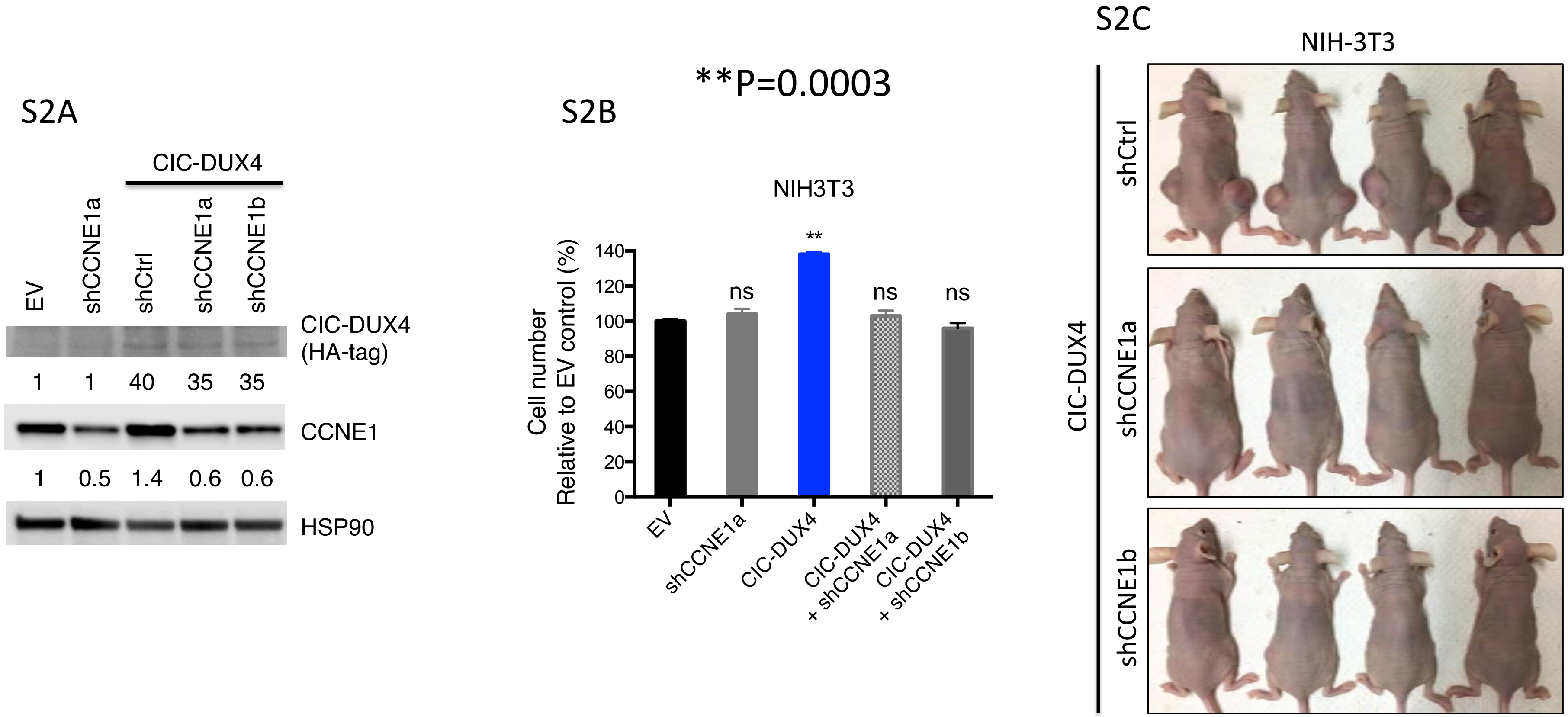

**Supplemental Figure 2. CCNE1 inhibition decreases tumor growth in CIC-DUX4 expressing cells.** A) Immunoblot of CIC-DUX4 (HA-Tag), CCNE1, and HSP90 in NIH-3T3 cells. B) Relative cell number of NIH-3T3 cells expressing either EV, shCCNE1a, CIC-DUX4 with or without shCCNE1a or shCCNE1b. **p-value = 0.0003. C) Subcutaneously implanted NIH-3T3 cells expressing CIC-DUX4 and either shCtrl, shCCNE1a, or shCCNE1b.

Supplemental Figure 3. Pharmacologic inhibition of CDK2 induces apoptosis in CIC-DUX4 expressing cells.

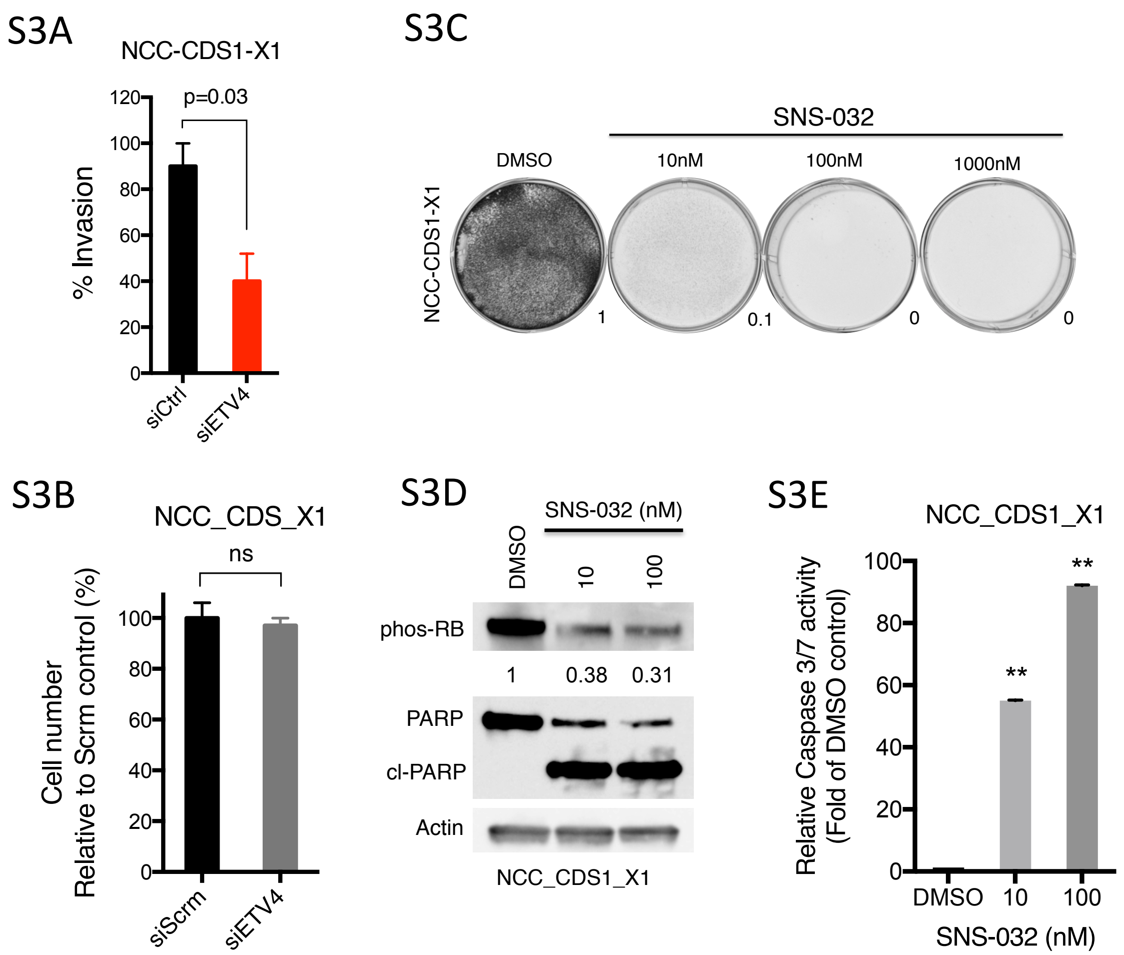

**Supplemental Figure 3. Pharmacologic inhibition of CDK2 induces apoptosis in CIC-DUX4 expressing cells.** A) Transwell invasion assay comparing CIC-DUX4 expressing NCC_CDS1_X1 cells with either siCtrl or siETV4. B) Relative cell number of NCC_CDS1_X1 cells following knockdown of ETV4 compared to scramble control. C) 72-hour crystal violet assay of NCC_CDS_X1 cells treated with SNS-032. D) immunoblot of phosphorylated-Rb, PARP, and Actin from NCC_CDS_X1 cells treated with SNS-032 or DMSO. E) Relative caspase 3/7 activity in NCC_CDS1_X1 cells treated with SNS-032 or DMSO. **p-value < 0.0001.

Supplemental Figure 4. The CCNE-CDK2 complex is a specific therapeutic target in CIC-DUX4 tumors.

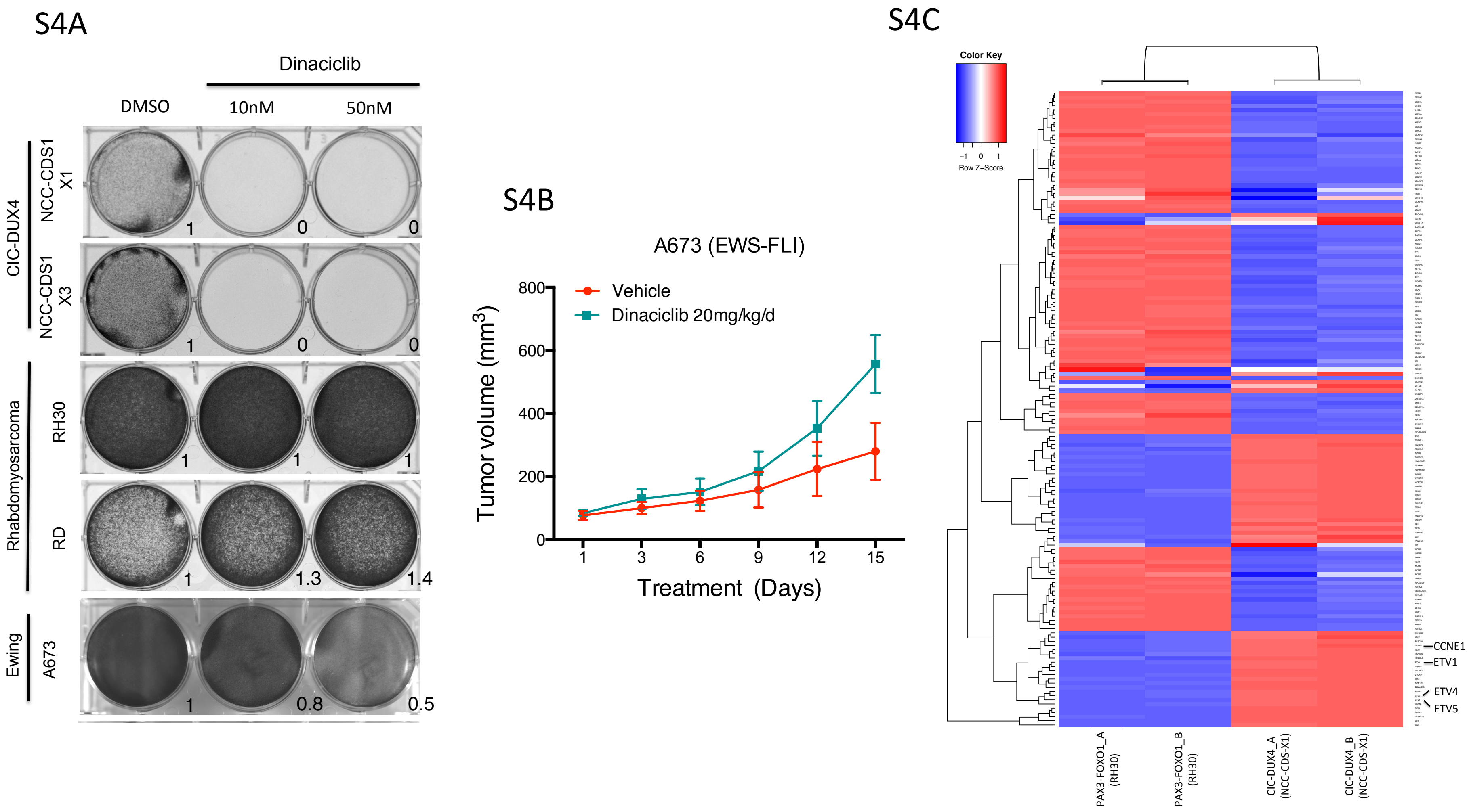

**Supplemental Figure 4. The CCNE-CDK2 complex is a specific therapeutic target in CIC-DUX4 tumors.** A) 72-hour crystal violet assay of CIC-DUX4 (NCC_CDS1_X1 and NCC_CDS_X3), rhabdomyosarcoma (RD and RH30), Ewing sarcoma (A673) cells treated with vehicle or dinaciclib. B) Subcutaneously implanted Ewing sarcoma (A673) cells treated with either vehicle or dinaciclib. C) Heatmap comparing 165 CIC-DUX4 activated genes identified in CIC-DUX4 expressing NCC-CDS1-X1 cells vs PAX3-FOXO1 containing RH30 cells. CCNE1, ETV1, ETV4, and ETV5 are magnified.

Supplemental Figure 5. Genetic inhibition of the CCNE-CDK2 complex decreases CIC-DUX4 tumor growth.

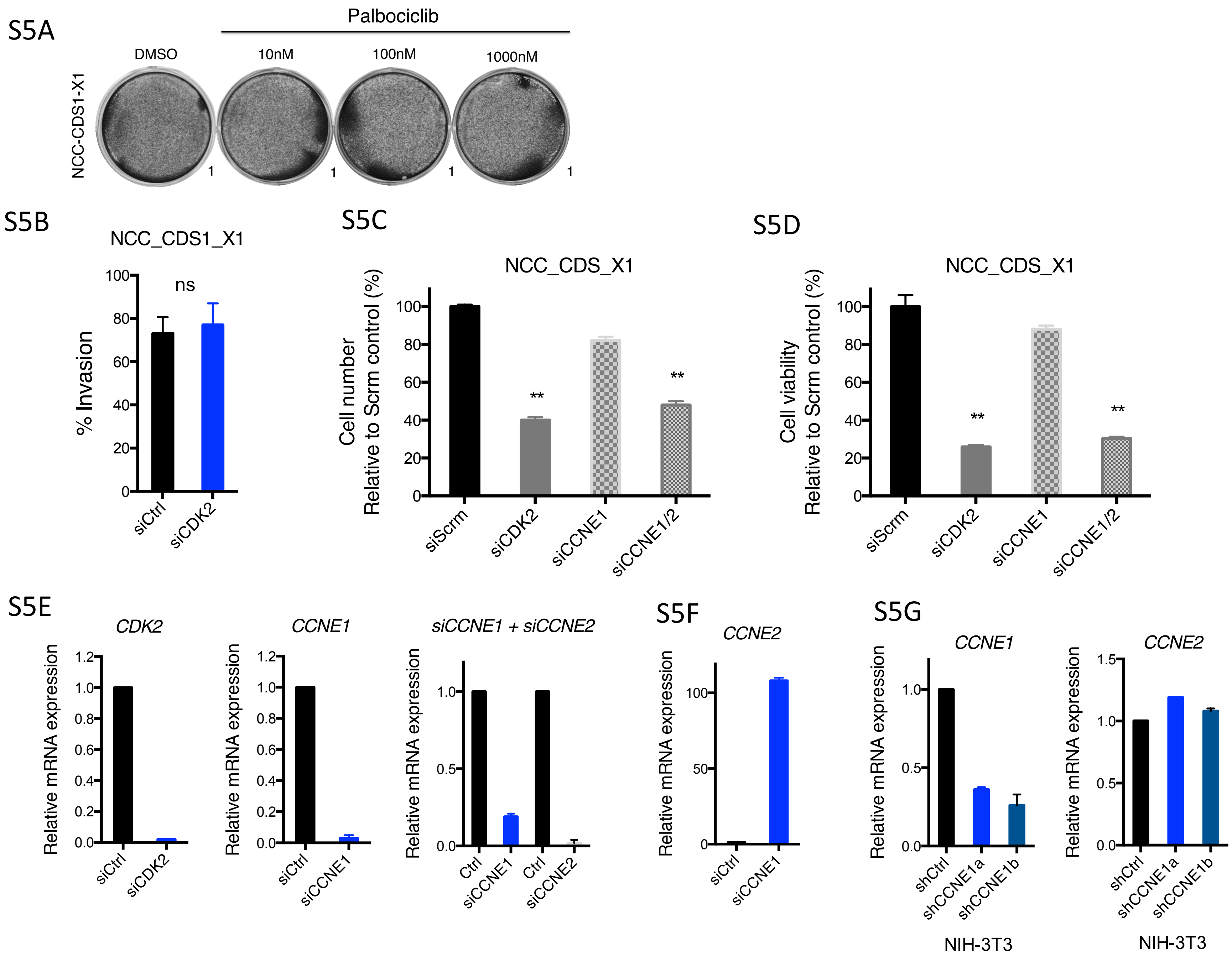

**Supplemental Figure 5. Genetic inhibition of the CCNE-CDK2 complex decreases CIC-DUX4 tumor growth.** A) Five-day crystal violet assay of NCC_CDS1_X1 cells treated with palbociclib. B) Transwell invasion assay comparing CIC-DUX4 expressing NCC_CDS1_X1 cells with either siCtrl or siCDK2. C) Relative cell number of NCC_CDS1_X1 cells following knockdown of CDK2, CCNE1, and combination CCNE1 and CCNE2 compared to scramble control. **p-value = 0.0001. D) Relative cell viability (cell titer glo assay) of NCC_CDS1_X1 cells following knockdown of *CDK2*, *CCNE1*, and combination *CCNE1* and *CCNE2* compared to scramble control. **p-value = 0.0001. E) Relative mRNA expression following *CDK2, CCNE1,* or dual *CCNE1* and *CCNE2* knockdown compared to scramble control. F) Relative CCNE2 mRNA expression following *CCNE1* knockdown compared to scramble control in NCC-CDS cells. G) Relative *CCNE1* mRNA expression following *CCNE1* knockdown compared to scramble control in NCC-CDS cells. H) Relative *CCNE2* mRNA expression following *CCNE1* knockdown compared to scramble control in CIC-DUX4 expressing NIH-3T3 cells.

| Table S1. 37 genes identified as putative CIC-DUX4 targets. | | |
|---|---|---|
| GeneSymbol | Description.x | ID.x |
| ANGPT2 | angiopoietin 2 | 285_at |
| CCNE1 | cyclin E1 | 898_at |
| CENPE | centromere protein E, 312kDa | 1062_at |
| CENPM | centromere protein M | 79019_at |
| CENPW | centromere protein W | 387103_at |
| CRH | corticotropin releasing hormone | 1392_at |
| CYP2S1 | cytochrome P450, family 2, subfamily S, polypeptide 1 | 29785_at |
| DLGAP5 | discs, large (Drosophila) homolog-associated protein 5 | 9787_at |
| EFR3B | EFR3 homolog B (S. cerevisiae) | 22979_at |
| ELOVL6 | ELOVL fatty acid elongase 6 | 79071_at |
| ETV4 | ets variant 4 | 2118_at |
| ETV5 | ets variant 5 | 2119_at |
| FAM83D | family with sequence similarity 83, member D | 81610_at |
| FANCI | Fanconi anemia, complementation group I | 55215_at |
| FLRT3 | fibronectin leucine rich transmembrane protein 3 | 23767_at |
| GLCCI1 | glucocorticoid induced transcript 1 | 113263_at |
| GTSE1 | G-2 and S-phase expressed 1 | 51512_at |
| HELLS | helicase, lymphoid-specific | 3070_at |
| HJURP | Holliday junction recognition protein | 55355_at |
| HMMR | hyaluronan-mediated motility receptor (RHAMM) | 3161_at |
| KIF4A | kinesin family member 4A | 24137_at |
| KIFC1 | kinesin family member C1 | 3833_at |
| MAD2L1 | MAD2 mitotic arrest deficient-like 1 (yeast) | 4085_at |
| MCM10 | minichromosome maintenance complex component 10 | 55388_at |
| MCM7 | minichromosome maintenance complex component 7 | 4176_at |
| POLQ | polymerase (DNA directed), theta | 10721_at |
| RAD51AP1 | RAD51 associated protein 1 | 10635_at |
| RHEBL1 | Ras homolog enriched in brain like 1 | 121268_at |
| RRM2 | ribonucleotide reductase M2 | 6241_at |
| SOSTDC1 | sclerostin domain containing 1 | 25928_at |
| SULT1E1 | sulfotransferase family 1E, estrogen-preferring, member 1 | 6783_at |
| SYBU | syntabulin (syntaxin-interacting) | 55638_at |
| TGFB3 | transforming growth factor, beta 3 | 7043_at |
| TGFBR3 | transforming growth factor, beta receptor III | 7049_at |
| VCAN | versican | 1462_at |
| VGF | VGF nerve growth factor inducible | 7425_at |
| ZWINT | ZW10 interactor, kinetochore protein | 11130_at |
